# Legionella felix sp. - A novel Legionella species isolated in Israel from hot tap water

**DOI:** 10.1101/2023.09.14.557692

**Authors:** Gal Zizelski Valenci, Shosanit Ohad, Mor Robinstein, Shereen Assaly, Ina Kutikov, Ludmila Groisman, Omer Murik, David A. Zeevi, Ravid Ben David, Alona Farber, Zeev Dveyrin, Efrat Rorman, Israel Nissan

## Abstract

Bacteria of the genus *Legionella* are natural pathogens of the environment that can cause legionellosis, which can result in severe human pneumonia called Legionnaires’ disease. Here we describe a novel *Legionella* species isolated from hot tap water. High quality complete genome was generated using a combination of Nanopore and Illumina sequencing approaches. Our novel *Legionella* isolate possesses a 3,913,681 bp chromosome, (GC content 38.78% Mol) and a single novel large *incF* conjugative plasmid of 160,500 bp (GC content 37.97%). Interestingly, the chromosome encodes for 14 secondary metabolites biosynthetic gene clusters, more than any known other *Legionella* sp. The novel plasmid encodes for multiple genes that confer resistance to heavy metals. Bioinformatics analysis including average nucleotide identity (ANI) and genome-based taxonomy computation revealed that the genome of the new isolate differs from any previously described *Legionella* species. The closest related species to our isolate is *Legionella cherrii*. The name proposed for the new specie is *Legionella felix* in honor of Dr. Arthur Felix (1887-1956), a pioneering microbiologist, and member of the Royal Society of UK, who established the National Public Health Laboratory in Tel Aviv.

## Introduction

*Legionella spp*. are Gram-negative bacteria associated with respiratory infections [1], which at the time of writing this article (April 2024), include 73 species, summarized in the “List of Prokaryotic names with Standing in Nomenclature” (www.bacterio.Net).

Bacteria of the *Legionella* genus are opportunistic human pathogens that can cause a serious type of pneumonia called Legionnaires’ disease and a mild flu-like illness, called Pontiac fever [2].

The most vulnerable population for Legionnaires’ disease are immunocompromised patients, including individuals treated with immunosuppressive therapy, chronic renal disease, age over 50 years, chronic cardiovascular disease, underlying respiratory disease, smoking, and diabetes [3, 4].

In nature, *Legionella spp*. are very common in the environment, populate fresh water and rarely cause illness. The bacteria parasitize and replicate within free-living eukaryotic phagotrophs, and protozoa species mainly *Amoebozoa* [1, 5].

According to the Centers for Disease Control and Prevention (CDC), *Legionella* can grow in natural as well as in non-natural settings, especially when not properly maintained.

*Legionella* can thrive in various water sources, including heated water, spa pools, air-conditioning cooling towers, premise plumbing, building water systems and geothermal water. *Legionella* persists in natural and man-made environments due to colonization of interspecies biofilms [2, 6]. Nevertheless, not much is known about *Legionella* in saline environments or seawater [7, 8].

*Legionella* infection occurs almost exclusively via breathing small droplets of water that contain the bacteria. In rare cases, while drinking goes mistakenly into the lungs, it can cause an infection (https://www.cdc.gov/legionella/downloads/fs-legionnaires.pdf). In general, person-to-person transmission is very rare [9].

In spite of the wide variety of *Legionella* species, only about 50% have been associated with human disease [10]. The species *L. pneumophila* is the cause for most Legionnaires’ disease (about 90%). *L. pneumophila* successfully adapted to new and challenging niches, such as Antarctic freshwater lakes, hot water sources (temperature over 60°C), and extremely acidic habitats [11-14]. The next most common etiological agents disease (2-7%) are; *L. longbeachae, L. bozemanii* and *L. micdadei* [12, 14]. The first detection of *L. cherrii* species in the respiratory samples of patients with pneumonia symptoms was published by *Atosa Ghorbani et al*., 2021 [15].

*Legionella* species encode a highly conserved type IVB secretion system (T4SS) called Dot/Icm. This system is essential for intracellular multiplication, and is encoded for about 29 different dot/icm genes [16]. The Dot/Icm system enables the bacteria to translocate effector proteins into the host cytosol, taking over cell functions and creating a specialized organelle, called the *Legionella* containing vacuole [17, 18].

In this work, we analyzed isolate #227, which belong to the *Legionella* spp. using genomic and phenotypic analysis, including high-throughput web server for genome-based prokaryote taxonomy - Type (Strain) Genome Server (TYGS) (https://tygs.dsmz.de) [19], microscopy and biochemical activity.

## Results and discussion

During a routine monitoring test of *Legionella* spp. from hot water spa, performed at the National Public Health Laboratory, Tel Aviv, isolate #227 showed special phenotypic characters. It was isolated from a hot water shower (tap water, temperature above 55°C). The sample was tested in accordance with standard procedures ISO 11731, for the isolation and quantitation of *Legionella* in water. This includes culture on Tryptic Soy Agar + Defibrinated Sheep Blood (blood agar, Hylabs, catalogue number PD049), and Buffered Charcoal Yeast Extract agar (BCYE, Hylabs, catalogue number PD073) at 36º C, for 48 hours in 2.5% CO2. No growth was observed on blood agar. Growth was observed in BCYE containing L-cysteine and iron, as anticipated from *Legionella* spp. We noticed that the colonies that grew on the selective plates were glowing under UV light (Figure 1). PCR analysis using the microproof® Legionella Quantification LyoKit (BIOTECON Diagnostics) identified it as *Legionella* sp.

**Figure 1.**
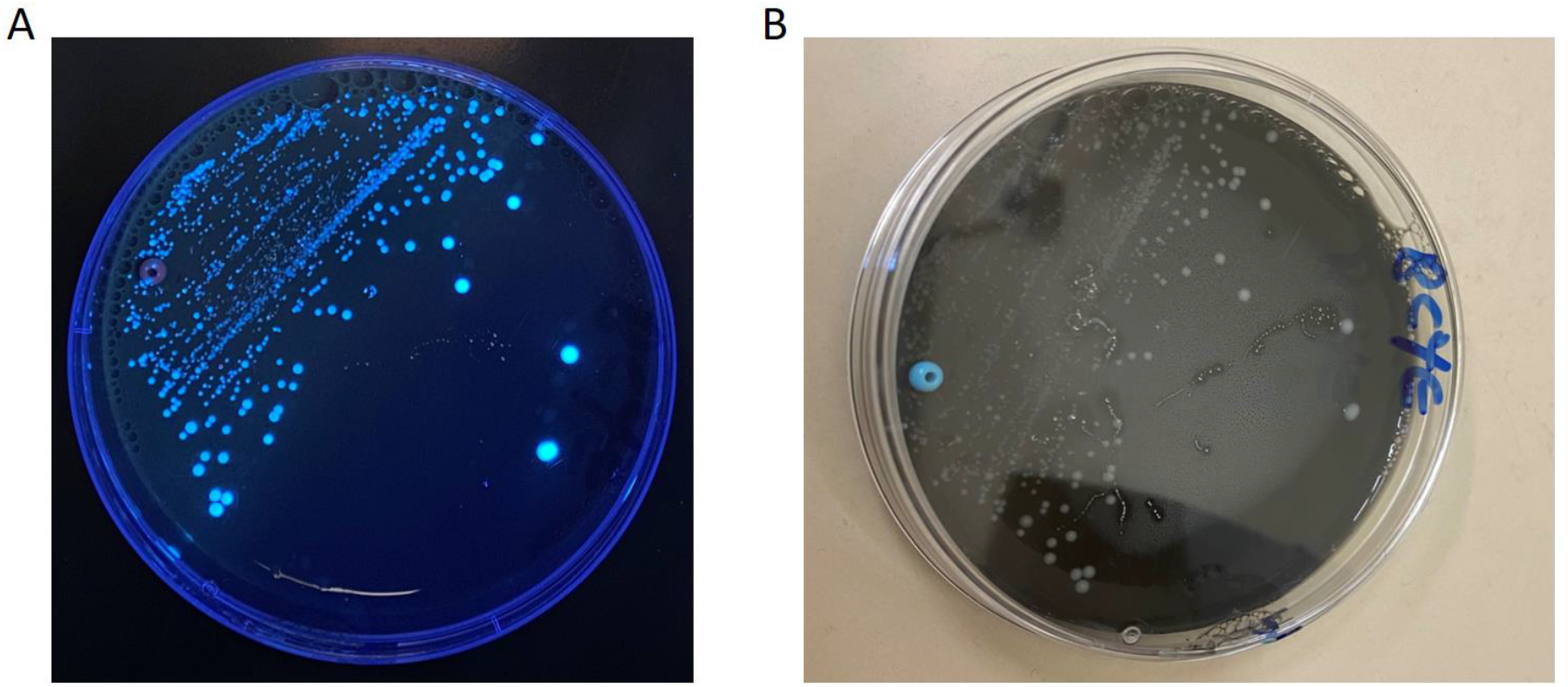
Growth of isolate 227 on Buffered Charcoal Yeast Extract agar (BCYE) at 36°C, for 48 hours in 2.5% CO2. Photo taken under UV (A) and visible light (B) respectively.

A matrix-assisted laser desorption ionization–time of flight mass spectrometry (MALDI-TOF MS) profile for isolate #227 was acquired on a Bruker Microflex system and interpreted with Bruker *Legionella* database using the MTB Compass, and Flex control software. Inactivation and preparation of the isolate for MALDI-TOF MS analysis were performed according to the manufacturer’s instructions. The MALDI-TOF-MS identified this isolate as *Legionella cherrii* at a maximal score of 2.03. Strain #227 appears in Gram stain as a large polymorphic Gram negative bacilli, with numerous elongated cells (figure 2). We tested the enzymatic profile of isolate #227 in comparison to *L. pneumophila* NCTC 11405 and *L. cherri* NCTC 11976, using the API® ZYM (bioMérieux), as described [20]. All tested strain generated identical biochemical profile (supplementary table number S1).

**Figure 2.**
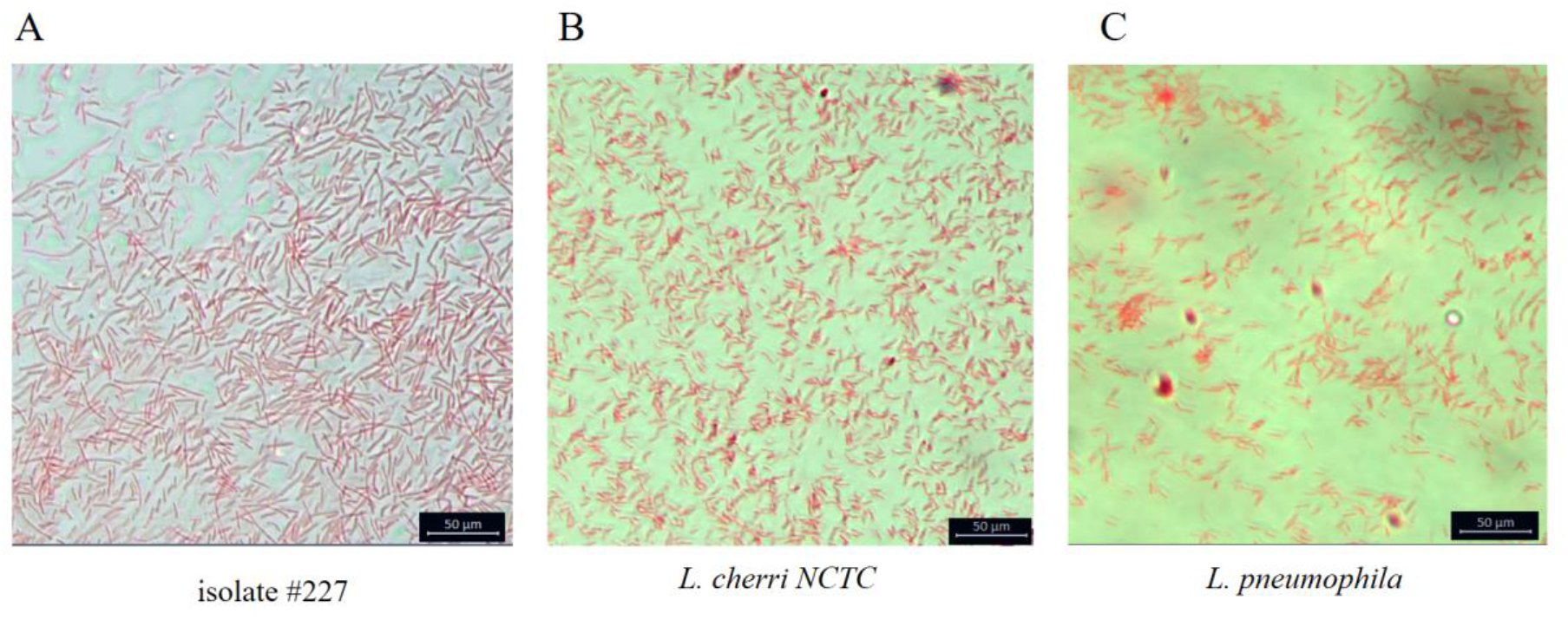
Image of Gram stain of isolate #227 (A), *L. cherri NCTC* (B), and *L. pneumophila NCTC* (C) under light microscope.

For Whole Genome Sequencing (WGS), 4 representative colonies were inoculated and suspended in 2 ml of Legionella - Pages Saline Buffer, followed by resuspension of 250 μl of bacterial suspension in 200 μl of Lysis buffer (Hylabs, catalogue number BP007). DNA was isolated using magLEAD® 12gC Automated Nucleic Acid Extraction System (Precision System Science Co., Ltd), following the manufacturer’s instructions. WGS of isolate #227 was carried out using two technologies as described; short-read sequencing on the Illumina MiSeq platform, and Long-read sequencing using the Oxford Nanopore platform. The sequences obtained from the two technologies were combined into two complete contigs (chromosome and plasmid). For the Illumina MiSeq platform, Paired-end library of isolate #227 was generated using Illumina DNA Prep library preparation kit according to Illumina protocols. The DNA library was sequenced using a 250-bp paired-end read MiSeq Reagent Kit v2 (500-cycles, catalog number MS-102-2003). All bioinformatics analyses were performed utilizing the Bacterial and Viral Bioinformatics Resource Center (BV-BRC) (https://www.bv-brc.org/) with default parameters, unless otherwise noted [21, 22]. The qualities of the raw reads were checked using Fastq Utilities of BV-BRC [22], confirming that the fastq files were of good quality.

For Oxford Nanopore (ONT) sequencing, DNA from isolate #227 was converted into a sequencing library using Rapid Barcoding Kit 96 (SQK-RBK110.96) chemistry according to the manufacturer’s protocol. Sequencing was performed on a MinION sequencer with an R9.4.1 (FLO-MIN106D) flow cell and base calling was carried out by Guppy version 6.0.1. Raw sequencing data were filtered with NanoFilt [23], keeping reads longer than 500bp and with quality > 10. Flye version 2.9 [24] was used for de novo assembly, and the assembly completeness was assessed with Quast version 5.2.0 and Busco version 5.4.2 [24-26]. The ONT sequences assembly results in two big contigs that serve as the template for Illumina reads mapping. A Consensus sequence was obtained with the BV-BRC Variation Analysis Service [22]. The schematic map of isolate #227 chromosome and plasmid sequence is presented in Figure 3. The assembled consensus genome was annotated using The BV-BRC Genome Annotation tool [22]. The genome comprises a 3,913,681 bp chromosome (GC content 38.78%) that encodes for 3505 CDS and a 160,500 bp large plasmid (GC content 37.97%) that encodes for 179 CDS (figure 3). Of the total 3,684 predicted protein-coding sequences, 1482 and 95 are hypothetical on the chromosome and plasmid respectively. A total of 7 rRNAs and 43 tRNAs were identified on the chromosome.

**Figure 3.**
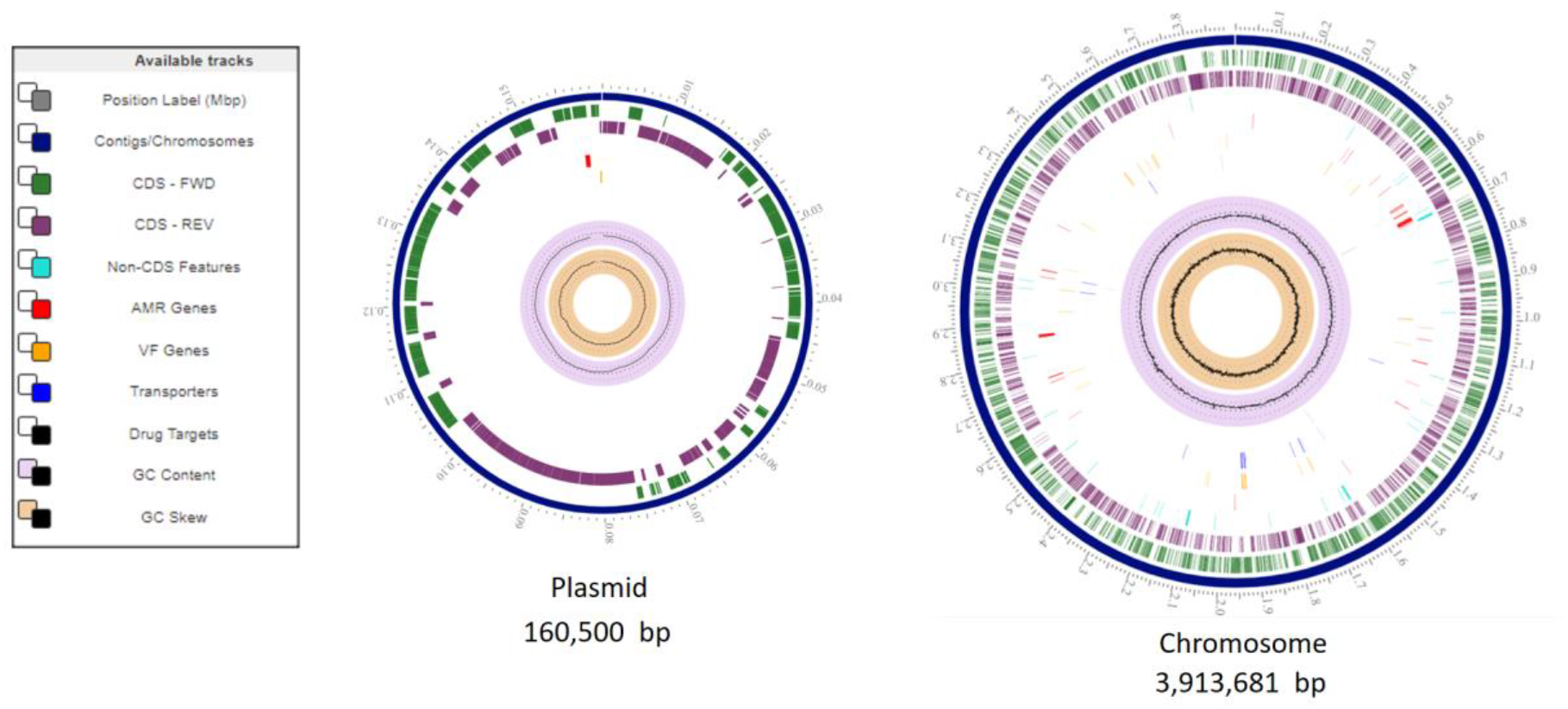
Schematic map of isolate #227 Chromosome and Plasmid. Isolate #227 consensus sequence genome was obtained by the BV-BRC Variation Analysis Service, combining Illumina and Nanopour technologies. ARM - Antimicrobial resistance, VF - virulence factor.

Examination of isolate #227 genome using the VFDB (Virulence Factor Database, as part of the BV-BRC), shows that the *Legionella* isolate #227 genome encodes for many virulence factors, including the dot/icm T4SS (supplementary table S2). Of the total plasmid genes, 93 encode for hypothetical proteins, 18 the *incF* plasmid conjugative transfer genes, 17 for heavy metal resistance, and few antibiotic resistance genes (supplementary table S3).

Notably, searching for plasmid homologues in the NCBI nr database, using Blastn revealed highly fragmented short homologues sequences mainly in other *Legionella sp*. plasmids, indicating that this is a novel plasmid.

We used the antibiotics and secondary metabolites analysis shell (antiSMASH 6.1.1) to search for the presence of secondary metabolite biosynthetic gene clusters in isolate #227 [27]. The analysis predicts 14 gene clusters encoding the biosynthesis of secondary metabolites and bioactive compounds (figure 4). For example, the antiSMASH analysis predicts four Non-ribosomal peptide synthetase cluster (NRPS) and NRPS-like region, one of them present 50% gene homology to Legioliulin biosynthetic gene cluster from *Legionella parisiensis* (figure 4, region 1.2). Legioliulin is a compound responsible for blue-white auto fluorescence in *Legionella dumoffii* under long-wavelength UV light [28]. Another example is the Betalactone biosynthesis coding region (figure 4, region 1.13), containing a gene of 13% sequence homology to a cluster of genes responsible for the synthesis of the Fengycin compound, a known antifungal lipopeptic antibiotic [29, 30]. Additionally, Type III polyketide synthases (PKS) region which contains genes with 12% & 25% homology to genes involved in the biosynthesis of Nematophin and Ambactin, respectively (figure 4, region 1.7). In the work of Nicolas J. Tobias et al., 2016 [31] he used antiSMASH for studying biosynthetic gene clusters (BGC) in 15 *Legionella* species. Indeed, *Legionella sp*. possesses variable BGCs, in particular 11 BGCs were identified in *L. anisa*. For the best of our knowledge, isolate #227 encodes for the largest known number of BGCs in a single *Legionella* genome. Notably, all these BGCs are located on the chromosome of isolate #227. The activities and potential utilizations of these BGCs could serve as a focus of a future dedicate study. The BV-BRC Similar Genome Finder Service, uses the Mash/MinHash algorithm, which presents each genome as a sketch, composed of 1000 of its k-mers [32].

**Figure 4.**
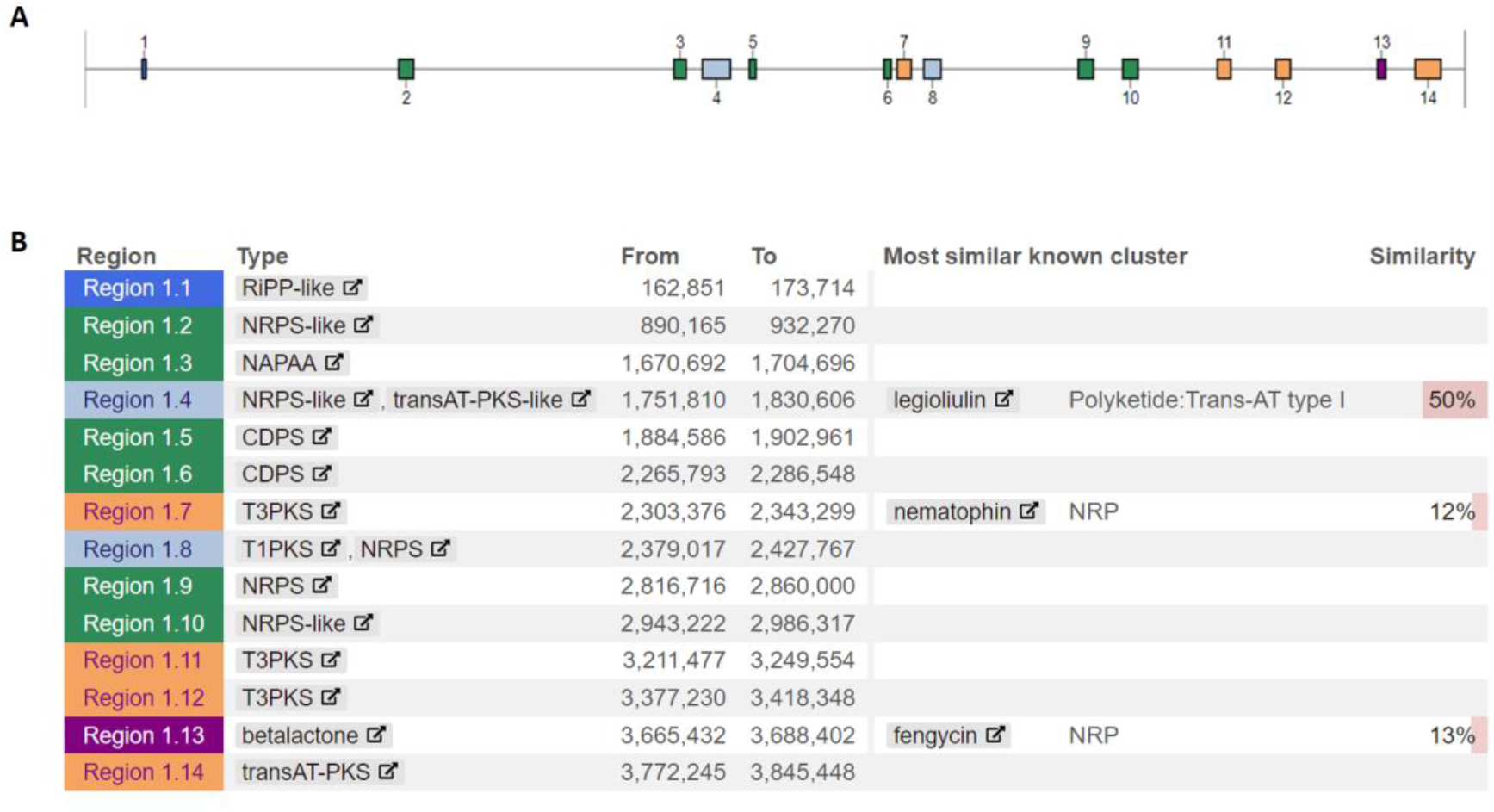
Prediction of gene clusters encoding the BGCs using antiSMASH 6.1.1. Schematic map of the chromosome with location of 14 BGCs (A) and list of regions prediction of gene clusters (B). The different color boxes in A, represents different clusters located on the schematic map of the genome.

This tool allows searching public genomic databases in the BV-BRC for the closest high-quality representative genome as determined by the NCBI, or, the closest homologue genome regardless of its quality of sequencing. As presented in supplementary table S4, the four most similar genomes strains in the database to isolate 227 are *L. cherrii*, that share between 193-203 common k-mers out of 1000. We used average nucleotide identity (ANI) calculator to estimate average nucleotide identity [33]. The level of similarity between the genome of isolate #227 and the most similar high quality representative genomes in the NCBI database (top 4 in supplementary table S4) is 93.9667, 93.6396, 93.902, 93.9381 respectively.

Genomes of the same species commonly show genome-aggregate ANI >95% among themselves [34] We performed genome-based taxonomy at TYPE (strain) GENOME SERVER (TYGS), according to the default parameters (https://tygs.dsmz.de/) [19, 35]. The TYGS is a well-known server connected to a large, continuously growing database of taxonomic, genomic, and nomenclatural information [19, 35-37].

The relevant type-strain genomes were determined automatically by the TYGS server. Genome-based phylogenetic trees of whole-genome and 16S rDNA gene are shown in figure The results of the taxonomic identification performed by the TYPE server are: “Potential new species detected. Isolate #227 does not belong to any species found in TYGS database”. Nevertheless, the most similar representative genome is *Legionella cherrii* DSM 19213 (Figure 5B). Notably, according to the 16S rDNA based tree (Figure 5A) *L. cherrii* is not the most similar to isolate #227, and the most similar species is *L. anisa*. Pairwise comparisons of isolate #227 genomes vs. type strain genomes and strains in dataset is available in supplementary Tables S5 and reference to the comparison genomes in supplementary Tables S6.

**Figure 5.**
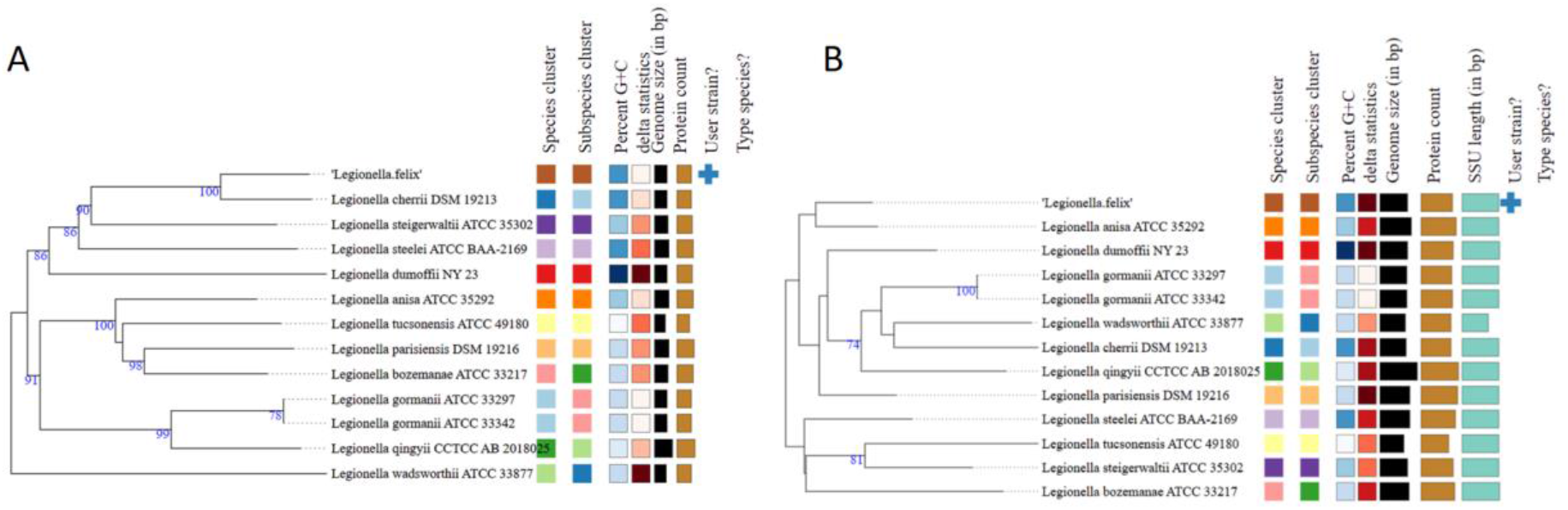
Whole-genome sequence-based phylogenetic tree (A) and 16S rDNA gene sequence-based tree (B). The trees represent the final step of the comprehensive genome-based taxonomic analyses available in the TYGS. Tree inferred with FastME 2.1.6.1. Leaf labels are annotated by affiliation to species and subspecies clusters, genomic G+C content, δ values, overall genome sequence length, number of proteins, and the kind of strain. Percent G+C 37.8-39.64, Delta statistics 0.299-0.449, Genome size (in bp) 3,573,650-5,227,415, SSU lengths (in bp) 1,092-1533. The colors of scheme boxes were determined automatically by the TYGS server.

Illumina and ONT sequences are available to the public at the NCBI (BioProject number PRJNA1004905).

Based on the above bioinformatics analysis we concluded that MLADI-TOF MS failed to correctly identify isolate #227.

In conclusion, by combining ONT and Illumina platforms, a complete and high quality sequence for isolate #227 was obtained. Based on meticulous bioinformatics analysis we found that *Legionella* isolate #277 represents a possible novel *Legionella* species that harbors large novel plasmid. Isolate #227 presents pleomorphic morphology that are different from its nearest relative *L. Cherrii*. We proposed the name *Legionella felix* for the new species [38, 39].

## References

1. Rowbotham, T.J., Preliminary report on the pathogenicity of Legionella pneumophila for freshwater and soil amoebae. J Clin Pathol, 1980. 33(12): p. 1179–83.

2. Chauhan, D. and S.R. Shames, Pathogenicity and Virulence of Legionella: Intracellular replication and host response. Virulence, 2021. 12(1): p. 1122–1144.

3. Cunha, B.A., A. Burillo, and E. Bouza, Legionnaires’ disease. Lancet, 2016. 387(10016): p. 376–385.

4. Barimani, M.J., Legionella: An uncommon cause of community-acquired pneumonia. JAAPA, 2022. 35(10): p. 38–42.

5. Swart, A.L., et al., Acanthamoeba and Dictyostelium as Cellular Models for Legionella Infection. Front Cell Infect Microbiol, 2018. 8: p. 61.

6. Rogers, J., et al., Influence of temperature and plumbing material selection on biofilm formation and growth of Legionella pneumophila in a model potable water system containing complex microbial flora. Appl Environ Microbiol, 1994. 60(5): p. 1585–92.

7. Gast, R.J., et al., Amoebae and Legionella pneumophila in saline environments. J Water Health, 2011. 9(1): p. 37–52.

8. Linsak, D.T., et al., Sea water whirlpool spa as a source of Legionella infection. J Water Health, 2021. 19(2): p. 242–253.

9. Borges, V., et al., Legionella pneumophila strain associated with the first evidence of person-to-person transmission of Legionnaires’ disease: a unique mosaic genetic backbone. Sci Rep, 2016. 6: p. 26261.

10. Fields, B.S., R.F. Benson, and R.E. Besser, Legionella and Legionnaires’ disease: 25 years of investigation. Clin Microbiol Rev, 2002. 15(3): p. 506–26.

11. McDade, J.E., et al., Legionnaires’ disease: isolation of a bacterium and demonstration of its role in other respiratory disease. N Engl J Med, 1977. 297(22): p. 1197–203.

12. Muder, R.R. and V.L. Yu, Infection due to Legionella species other than L. pneumophila. Clin Infect Dis, 2002. 35(8): p. 990–8.

13. Oliva, G., T. Sahr, and C. Buchrieser, The Life Cycle of L. pneumophila: Cellular Differentiation Is Linked to Virulence and Metabolism. Front Cell Infect Microbiol, 2018. 8: p. 3.

14. Salloum, G., et al., Identification of Legionella species by ribotyping and other molecular methods. Res Microbiol, 2002. 153(10): p. 679–86.

15. Ghorbani, A., et al., Occurrence of the Legionella species in the respiratory samples of patients with pneumonia symptoms from Ahvaz, Iran; first detection of Legionella cherrii. Mol Biol Rep, 2021. 48(11): p. 7141–7146.

16. Burstein, D., et al., Genomic analysis of 38 Legionella species identifies large and diverse effector repertoires. Nat Genet, 2016. 48(2): p. 167–75.

17. Qiu, J. and Z.Q. Luo, Legionella and Coxiella effectors: strength in diversity and activity. Nat Rev Microbiol, 2017. 15(10): p. 591–605.

18. Mace, K., et al., Proteins DotY and DotZ modulate the dynamics and localization of the type IVB coupling complex of Legionella pneumophila. Mol Microbiol, 2022. 117(2): p. 307–319.

19. Meier-Kolthoff, J.P., et al., TYGS and LPSN: a database tandem for fast and reliable genome-based classification and nomenclature of prokaryotes. Nucleic Acids Res, 2022. 50(D1): p. D801–D807.

20. Muller, H.E., Enzymatic profile of Legionella pneumophilia. J Clin Microbiol, 1981. 13(3): p. 423–6.

21. Wattam, A.R., et al., Improvements to PATRIC, the all-bacterial Bioinformatics Database and Analysis Resource Center. Nucleic Acids Res, 2017. 45(D1): p. D535–D542.

22. Olson, R.D., et al., Introducing the Bacterial and Viral Bioinformatics Resource Center (BV-BRC): a resource combining PATRIC, IRD and ViPR. Nucleic Acids Res, 2022.

23. De Coster, W., et al., NanoPack: visualizing and processing long-read sequencing data. Bioinformatics, 2018. 34(15): p. 2666–2669.

24. Mikheenko, A., et al., Versatile genome assembly evaluation with QUAST-LG. Bioinformatics, 2018. 34(13): p. i142–i150.

25. Seppey, M., M. Manni, and E.M. Zdobnov, BUSCO: Assessing Genome Assembly and Annotation Completeness. Methods Mol Biol, 2019. 1962: p. 227–245.

26. Simao, F.A., et al., BUSCO: assessing genome assembly and annotation completeness with single-copy orthologs. Bioinformatics, 2015. 31(19): p. 3210–2.

27. Blin, K., et al., antiSMASH 6.0: improving cluster detection and comparison capabilities. Nucleic Acids Res, 2021. 49(W1): p. W29–W35.

28. Amemura-Maekawa, J., et al., Legioliulin, a new isocoumarin compound responsible for blue-white autofluorescence in Legionella (Fluoribacter) dumoffii under long-wavelength UV light. Biochem Biophys Res Commun, 2004. 323(3): p. 954–9.

29. Vanittanakom, N., et al., Fengycin--a novel antifungal lipopeptide antibiotic produced by Bacillus subtilis F-29-3. J Antibiot (Tokyo), 1986. 39(7): p. 888–901.

30. Sur, S., T.D. Romo, and A. Grossfield, Selectivity and Mechanism of Fengycin, an Antimicrobial Lipopeptide, from Molecular Dynamics. J Phys Chem B, 2018. 122(8): p. 2219–2226.

31. Tobias, N.J., et al., Legionella shows a diverse secondary metabolism dependent on a broad spectrum Sfp-type phosphopantetheinyl transferase. PeerJ, 2016. 4: p. e2720.

32. Ondov, B.D., et al., Mash: fast genome and metagenome distance estimation using MinHash. Genome Biol, 2016. 17(1): p. 132.

33. Goris, J., et al., DNA-DNA hybridization values and their relationship to whole-genome sequence similarities. Int J Syst Evol Microbiol, 2007. 57(Pt 1): p. 81–91.

34. Rodriguez, R.L., et al., An ANI gap within bacterial species that advances the definitions of intra-species units. mBio, 2024. 15(1): p. e0269623.

35. Meier-Kolthoff, J.P. and M. Goker, TYGS is an automated high-throughput platform for state-of-the-art genome-based taxonomy. Nat Commun, 2019. 10(1): p. 2182.

36. de Oliveira Sant’Anna, L., et al., Corynebacterium guaraldiae sp. nov.: a new species of Corynebacterium from human infections. Braz J Microbiol, 2023. 54(2): p. 779–790.

37. Prescott, R.D., et al., Bridging Place-Based Astrobiology Education with Genomics, Including Descriptions of Three Novel Bacterial Species Isolated from Mars Analog Sites of Cultural Relevance. Astrobiology, 2023. 23(12): p. 1348–1367.

38. Craigie, J., ARTHUR FELIX 1887-1956. Royal Society (United Kingdom).

39. Nicolle, P., [Arthur Felix; 1887-1956]. Bull Soc Pathol Exot Filiales, 1956. 49(2): p. 232–5.

